# The role of liver fat in cardiometabolic diseases is highlighted by genome-wide association study of MRI-derived measures of body composition

**DOI:** 10.1101/2022.02.24.481887

**Authors:** Dennis van der Meer, Tiril P. Gurholt, Ida E. Sønderby, Alexey A. Shadrin, Guy Hindley, Zillur Rahman, Ann-Marie G. de Lange, Oleksandr Frei, Olof D. Leinhard, Jennifer Linge, Rozalyn Simon, Lars T. Westlye, Sigrun Halvorsen, Anders M. Dale, Tom H. Karlsen, Tobias Kaufmann, Ole A. Andreassen

## Abstract

**Background & Aims:** Obesity and associated morbidities, metabolic associated liver disease (MAFLD) included, constitute some of the largest public health threats worldwide. Body composition and related risk factors are known to be heritable and identification of their genetic determinants may aid in the development of better prevention and treatment strategies. Recently, large-scale whole-body MRI data has become available, providing more specific measures of body composition than anthropometrics such as body mass index. Here, we aimed to elucidate the genetic architecture of body composition, by conducting the first genome-wide association study (GWAS) of these MRI-derived measures.

**Methods:** We ran both univariate and multivariate GWAS on fourteen MRI-derived measurements of adipose and muscle tissue distribution, derived from scans from 34,036 White European UK Biobank participants (mean age of 64.5 years, 51.5% female).

**Results:** Through multivariate analysis, we discovered 108 loci with distributed effects across the body composition measures and 256 significant genes primarily involved in immune system functioning. Liver fat stood out, with a highly discoverable and oligogenic architecture and the strongest genetic associations. Comparison with 21 common cardiometabolic traits revealed both shared and specific genetic influences, with higher mean heritability for the MRI measures (h^2^=.25 vs. .16, p=1.4×10^−6^). We found substantial genetic correlations between the body composition measures and a range of cardiometabolic diseases, with the strongest correlation between liver fat and type 2 diabetes (r_g_=.48, p=1.6×10^−22^).

**Conclusions:** These findings show that MRI-derived body composition measures complement conventional body anthropometrics and other biomarkers of cardiometabolic health, highlighting the central role of liver fat, and improving our knowledge of the genetic architecture of body composition and related diseases.

## Introduction

Obesity and associated cardiometabolic diseases are currently considered one of the largest global public health concerns^1,2^. Over one-third of the United States adult population qualifies for a diagnosis of metabolic syndrome,^3^ characterized by excessive visceral adiposity, insulin resistance, hypertension, low high-density lipoprotein cholesterol, and hypertriglyceridemia.^4,5^ Metabolic syndrome substantially increases the risk of coronary artery disease, type 2 diabetes, cancer, and metabolic associated fatty liver disease (MAFLD, previously described as non-alcoholic fatty liver disease^6^).^7–11^ Body composition is also associated with brain structure and brain disorders.^12,13^ An improved understanding of the genetic and biological determinants of body composition is needed to provide insights into the complex interplay between metabolic factors, prevent and treat multiple highly prevalent conditions, and improve public health outcomes.^2,10^

Body composition is partly determined by a complex constellation of interacting metabolic processes and inter-organ cross-talk that may become dysregulated and lead to metabolic syndrome.^14^ In susceptible individuals, excessive energy intake, stored as visceral adipose tissue, combined with insulin resistance, leads to heightened lipolysis and release of free fatty acids.^15^ Increased free fatty acid flux to the liver results in hypertriglyceridemia, which in turn contributes to dyslipidemia and atherosclerosis. Lipolysis in visceral adipose tissue further promotes insulin resistance and gluconeogenesis and increases pro-inflammatory reactions that exacerbate endothelial dysfunction and hypertension.^15^ This is reflected in heightened levels of pro-inflammatory markers among individuals with metabolic syndrome.^16^ Muscle mass is also a determinant of cardiometabolic health,^17^ as skeletal muscle constitutes the largest insulin-sensitive tissue in the body and is the primary site for insulin-stimulated glucose utilization.^18^ Still, the nature and extent of overlap between these different determinants of cardiometabolic functioning remain unclear.

Measures of localized adipose tissue, liver fat and regional muscle volume can now be accurately extracted from whole-body MRI scans.^19–22^ MRI-based body tissue quantification offers more sensitive proxies of cardiometabolic health than body anthropometrics such as waist circumference and body mass index (BMI)^23^, which also lack a direct connection to pathophysiology.^5,24^ Measures of regional adipose tissue, most accurately and comprehensively identified through MRI,^25,26^ show independent associations with cardiometabolic diseases and improve risk prediction beyond body anthropometrics.^27–29^

In addition to social and physical environmental factors,^30^ genetically determined individual differences play a significant role in regulating body composition.^31–33^ Cardiometabolic risk factors have both unique and shared genetic correlates.^34^ Much less is known about the genetics of specific MRI-derived body composition measures.^35^ We aimed to map the unique and shared genetic architectures across the MRI-derived body composition to provide a holistic understanding of the interplay between different tissue types and their role in metabolic syndrome and cardiometabolic health.

## Results

We conducted GWASs of fourteen MRI-derived muscle and adipose tissue distribution measures and investigated the genetic link to conventional cardiometabolic risk factors. We included six measures of adipose tissue distribution: abdominal subcutaneous adipose tissue, visceral adipose tissue, abdominal fat ratio, anterior and posterior thigh muscle fat infiltration, and liver protein density fat fraction. Additionally, we investigated three measures related to thigh muscle tissue, namely anterior and posterior thigh muscle volume and weight-to-muscle ratio. We further analyzed visceral and abdominal adipose tissue, and anterior and posterior muscle volume, divided by standing height in meters squared, and total thigh muscle volume z-score (sex-, height-, weight- and BMI-invariant).^36^ See Table 1 for an overview of these measures, and the Methods section for protocols and definitions. Given a total of fourteen individual measures, we set the univariate GWAS significance threshold at α=5*10^−8^/14=3.6*10^−9^. Our sample for the main analyses consisted of 34,036 White European participants of the UK Biobank (UKB), with a mean age of 64.5 years (standard deviation (SD) 7.4 years), 51.5% female. We pre-residualized all measures for age, sex, test center, and the first twenty genetic principal components to control for population stratification.

**Table 1.** MRI-derived measures of body composition included in this study, together with the available sample size and number of loci discovered through univariate GWAS.

| *Measure* | *Abbreviation* | *N* | *# loci* |
| --- | --- | --- | --- |
| Abdominal subcutaneous adipose tissue | ASAT | 33979 | 1 |
| Visceral adipose tissue | VAT | 33989 | 2 |
| Anterior thigh muscle volume | ATMV | 33415 | 7 |
| Posterior thigh muscle volume | PTMV | 33459 | 10 |
| Anterior thigh muscle fat infiltration (%) | ATMFI | 33347 | 19 |
| Posterior thigh muscle fat infiltration (%) | PTMFI | 33392 | 25 |
| Weight-muscle-ratio | WMR | 33406 | 1 |
| Abdominal fat ratio | AFR | 33375 | 1 |
| Liver proton density fat fraction (%) | LPDFF | 33674 | 8 |
| VAT/height<sup>2</sup> | VATi | 33007 | 2 |
| ASAT/height<sup>2</sup> | ASATi | 32998 | 1 |
| ATMV/height<sup>2</sup> | ATMVi | 32442 | 5 |
| PTMV/height<sup>2</sup> | PTMVi | 32483 | 0 |
| Total thigh muscle volume z-score | TTMVz | 32401 | 5 |

### Univariate GWAS

Univariate GWASs on the individual measures revealed a total of 87 loci, including 54 unique, surpassing the study-wide significance threshold of 3.6*10^−9^. Two loci stood out with highly significant p-values, on chromosome 19 (lead rs58542926, p=8.8×10^−113^) and chromosome 22 (lead rs738409, p=4.6×10^−166^), both identified in the GWAS on liver fat. Using converging positional, eQTL, and chromatin interaction information (see Methods), we mapped these loci to genes previously coupled to MAFLD (rs738409: *PNPLA3, SAMM50, PARVB*)^37^ as well as inflammatory processes and cancer (rs58542926: *CD99*).^38^ The Supplementary Information (SI) contains Manhattan plots and overviews of all loci discovered together with mapped genes.

Additionally, we assessed the generalization of the discovered loci in a hold-out set of 5,081 non-White European UKB participants with identical processing steps. Of the 84 lead SNPs available in this set, 82 had effects in the same direction as the main analyses (97.6%, sign-test p<1×10^−16^). Thus, our results suggest a cross-ethnicity generalization of these genetic associations with MRI-derived measures of body composition, despite the known high variability of body anthropometrics across ethnicities.^5,24^

In total, we identified eight study-wide significant loci for liver fat, validating those found in a previous smaller GWAS.^35^ Gene-based analysis through Multi-marker Analysis of GenoMic Annotation (MAGMA) identified 35 genome-wide significant genes, including the three primary MAFLD genes (*TM6SF2* p=7.2*10^−16^, *PNPLA3* p=1.0*10^−14^, and *TMC4-MBOAT7* p=2.1*10^−8^),^39– 42^ further confirming the strong biological validity of this liver fat measure and its connection to MAFLD. Functional annotation of the set of 35 genes revealed differential expression in the liver, pancreas, and subcortical brain regions and significant enrichment among Gene Ontology (GO) biological processes specifically related to lipid homeostasis and metabolic processes. The SI further contains results of gene set enrichment analyses for each individual measure.

Next, we estimated the polygenicity and effect size variance (‘discoverability’) by fitting a Gaussian mixture model of null and non-null effects to the GWAS summary statistics using MiXeR.^43,44^ The results are summarized in Figure 1a, depicting the estimated proportion of genetic variance explained by discovered SNPs for each measure as a function of sample size. This illustrates that body MRI measures generally show genetic architectures similar to e.g. brain MRI measures, characterized by high polygenicity.^45,46^ However, the notable exception is liver fat, with substantially lower polygenicity and higher discoverability than the other measures, in line with the relatively few highly significant associations we identified through the GWAS.

**Figure 1.**
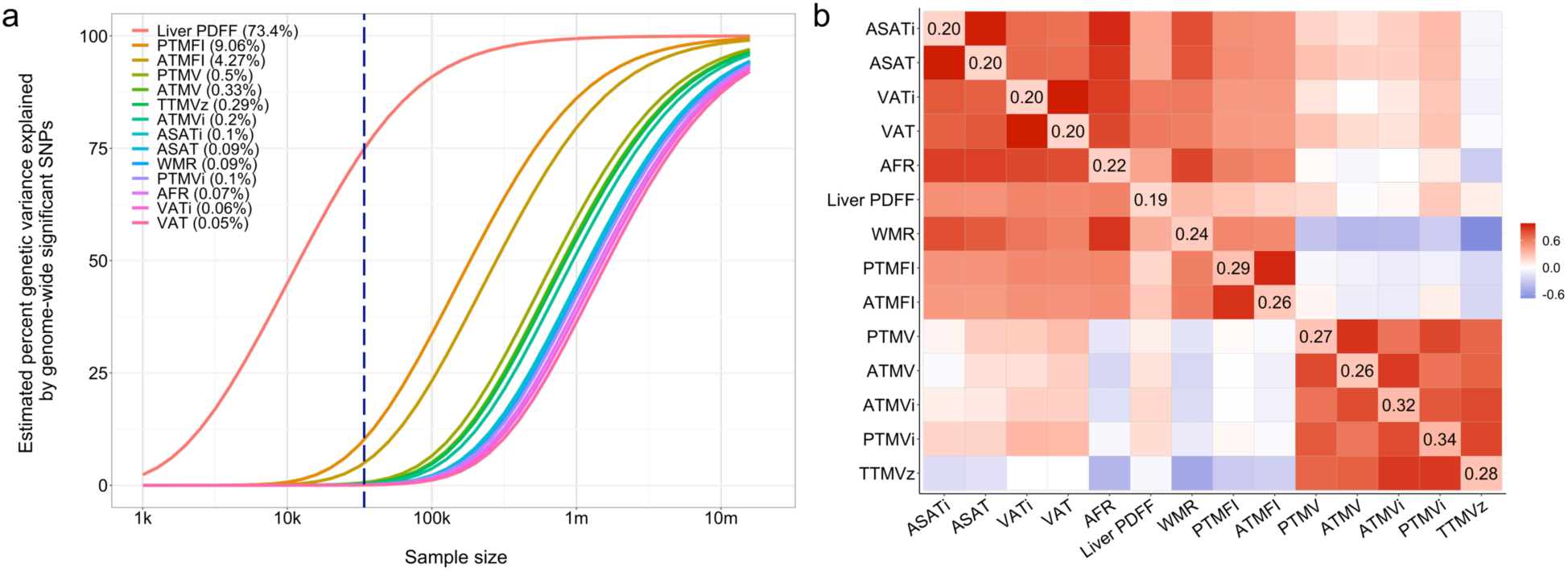
Comparison of the genetic architecture of individual body composition measures. **a)** The relation between genetic variance explained by genome-wide significant hits (y-axis) and sample size (x-axis) for each measure (solid colored lines). The vertical dashed blue line marks the current sample size, with the corresponding percent genetic variance explained indicated between brackets in the legend. **b)** Correlation between the measures, with phenotypic correlation shown in bottom triangle and genetic correlation in the upper triangle, and heritability on the diagonal. Abbreviations: ASAT=abdominal subcutaneous adipose tissue, VAT=visceral adipose tissue, AFR= abdominal fat ratio, WMR=weight-muscle-ratio, ATMV=anterior thigh muscle volume, PTMV=Posterior thigh muscle volume, ATMFI=anterior thigh muscle fat infiltration, PTMFI=posterior thigh muscle fat infiltration, Liver PDFF=liver proton density fat fraction, TTMVz=total thigh muscle volume z-score, i=index, referring to a measure divided by standing height^2^.

Figure 1b visualizes the phenotypic and genetic correlations between each pair of measures, confirming a strong structure and a subdivision between adipose- and muscle-related measures. SNP-based heritability ranged from 19% to 34% (all p<1×10^−16^); see the diagonal of Figure 1b.

### Multivariate GWAS

Gene variants are likely to have distributed effects across these measures, as they are correlated components of the same biological system. We therefore also jointly analyzed all measures through the Multivariate Omnibus Statistical Test (MOSTest),^47^ which increases statistical power in a scenario of shared genetic signal across the univariate measures.^47–49^ After applying a rank-based inverse normal transformation, we performed MOSTest on the residualized measures, yielding a multivariate association with 9.1 million SNPs included.

MOSTest revealed 108 significant independent loci across all MRI-derived measures (see Figure 2a and SI). Figure 2b visualizes the significance of the association between the individual measures and each of the 108 loci, illustrating the presence of many shared but also specific genetic variants.

**Figure 2.**
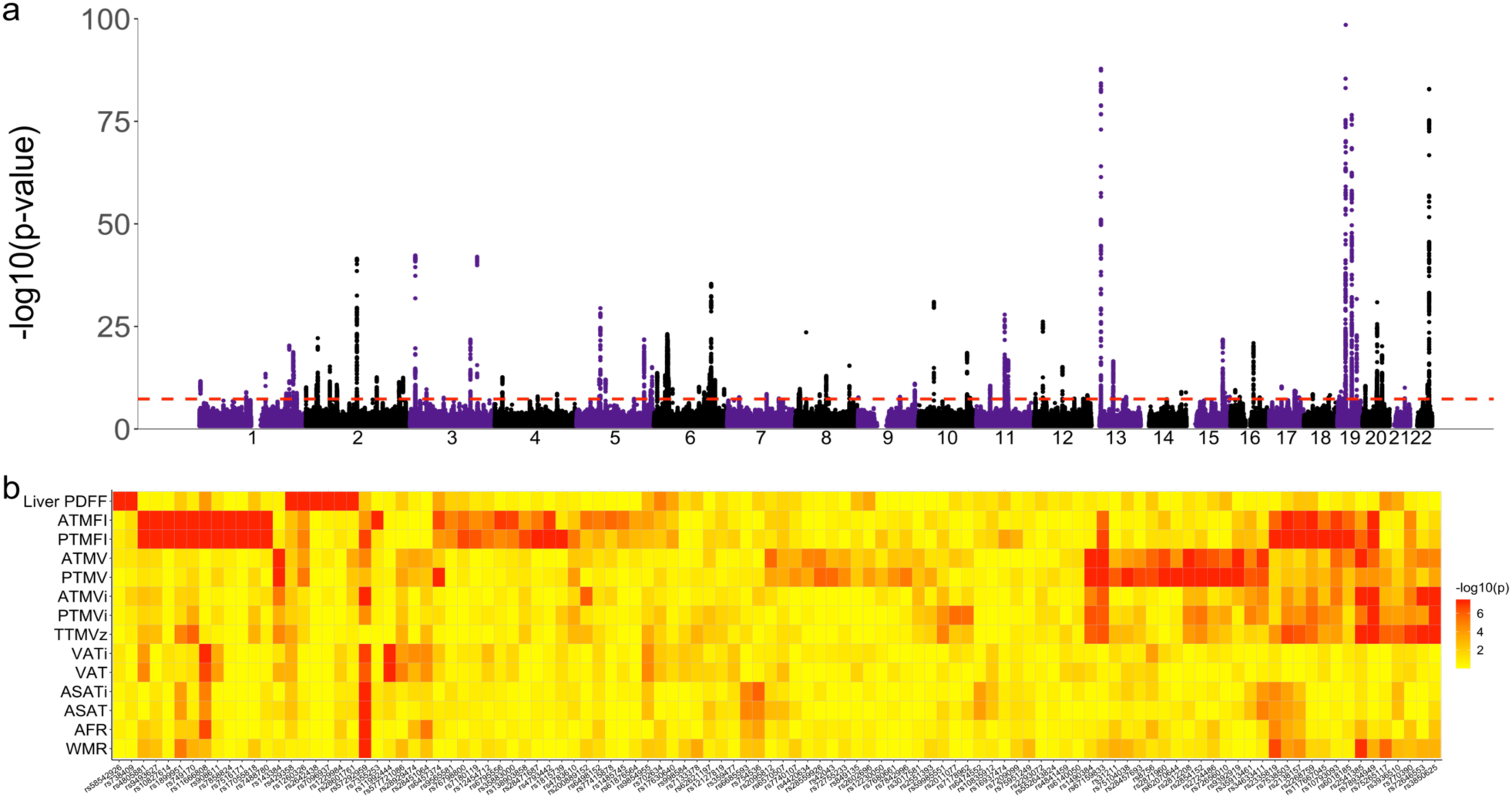
Multivariate locus discovery. **a)** Manhattan plot of the multivariate GWAS on all MRI-derived body composition measures, with the observed -log10(p) of each SNP shown on the y-axis. The x-axis shows the relative genomic location, grouped by chromosome, and the red dashed line indicates the whole-genome significance threshold of 5×10^−8^. The y-axis is clipped at -log10(p)=75. **b)** Heatmap showing -log10(p) of the association between the lead variants of MOSTest-identified independent loci (x-axis) and each of the individual MRI measures (y-axis). The values are capped at 7.5 (p=5×10^−8^).

MAGMA identified 256 significant genes after multiple comparison correction (α=.05/18,203), with highly significant differential expression in the liver, pancreas, heart, muscle, and several other tissues (Figure 3). Coupling the significant genes to the Reactome database^50^ indicated most prominent associations with the adaptive immune system and cytokine signaling (p<1*10^−16^), see Supplementary Data and Supplementary Figure 2 for an overview.

**Figure 3.**
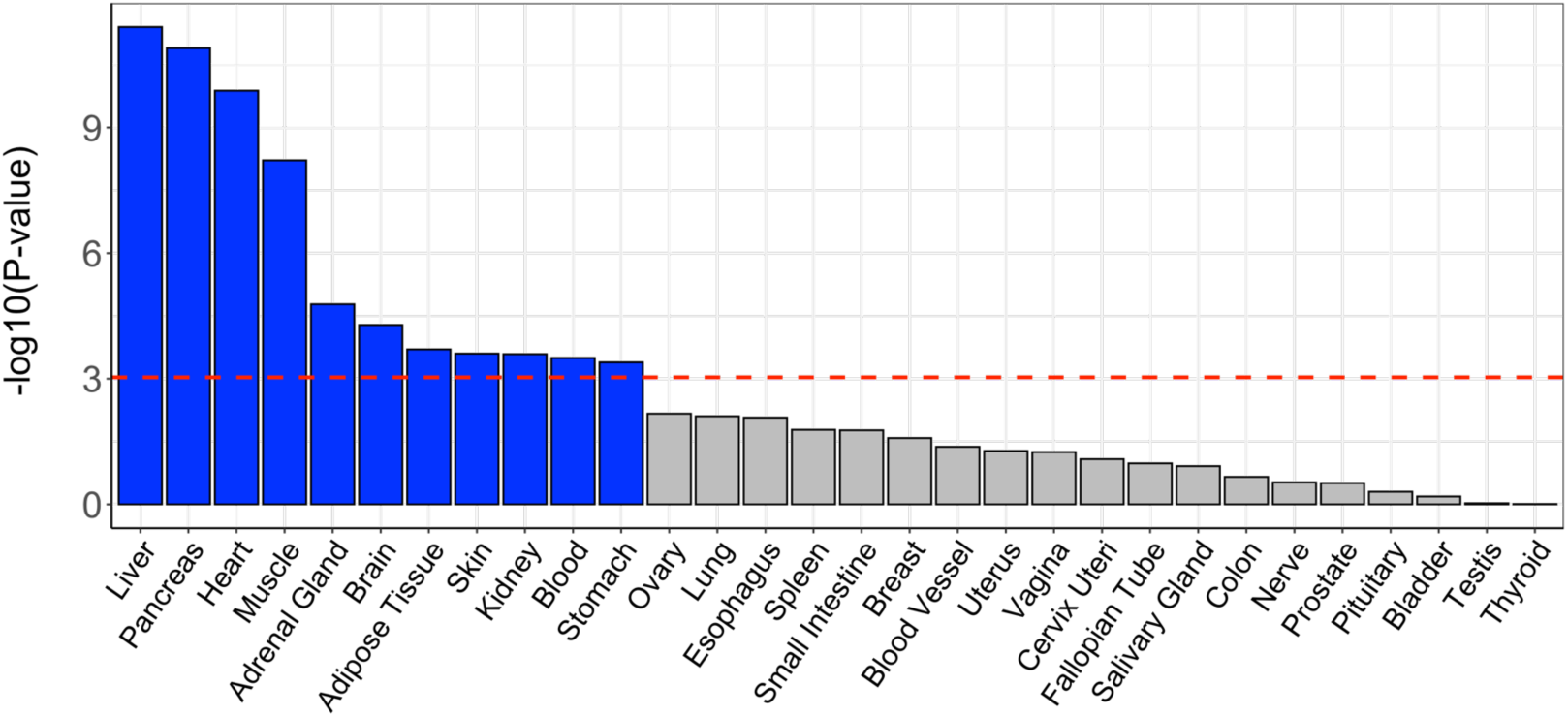
Tissue-specific differential expression of the set of significant genes identified through the multivariate GWAS on MRI-derived measures of body composition. The red-dotted line indicates the multiple comparisons-corrected significance threshold.

### Comparison of genetic architecture with cardiometabolic risk factors

We additionally analyzed a set of 21 measures of anthropometric and cardiometabolic factors (e.g., BMI, triglycerides, cholesterol, blood pressure; see Table 2), which were available for up to 412,316 White European UKB participants. Through multivariate GWAS on this separate set of measures in the full UKB sample, we found 1134 genome-wide significant loci with α=5×10^−8^ (list provided in SI). Of the 108 loci identified through the primary multivariate analysis of MRI-derived body composition measures, 94 (87%) were significant in this secondary analysis in a larger sample. This indicates that these sets of measures overall are influenced by the same network of biological processes.

**Table 2.** Measures of cardiometabolic health used in the secondary analyses, together with abbreviations and available sample sizes.

| *Measure* | *Abbreviation* | *N* |
| --- | --- | --- |
| Cholesterol |  | 389832 |
| High-density lipoproteins | HDL | 356802 |
| Low-density lipoproteins | LDL | 389115 |
| Triglycerides |  | 389524 |
| Apolipoprotein A |  | 354829 |
| Apolipoprotein B |  | 387945 |
| Cholesterol to HDL |  | 356734 |
| ApoB to ApoA |  | 353056 |
| C-Reactive Protein | CRP | 388993 |
| Glucose |  | 356558 |
| Glycated hemoglobin | HbA1c | 389728 |
| Alanine aminotransferase | ALT | 389701 |
| Aspartate aminotransferase | AST | 388406 |
| Gamma-glutamyl transferase | GGT | 389634 |
| Creatinine |  | 389637 |
| Body mass index | BMI | 407558 |
| Waist circumference |  | 408179 |
| Hip circumference |  | 408134 |
| Waist-to-hip ratio | WHR | 408096 |
| Diastolic blood pressure | DBP | 36591 |
| Systolic blood pressure | SBP | 36591 |

The heritability of the MRI-derived measures (mean h^2^=.25) was significantly higher than the body anthropometrics and other biomarkers (mean h^2^=.16), p=1.4×10^−6^. As shown in Figure 4, these measures generally showed higher genetic correlations with the MRI-derived measures of adipose tissue than the muscle-related measures. Further, BMI, hip/waist circumference, and waist-to-hip-ratio were genetically correlated with nearly all body MRI measures.

**Figure 4.**
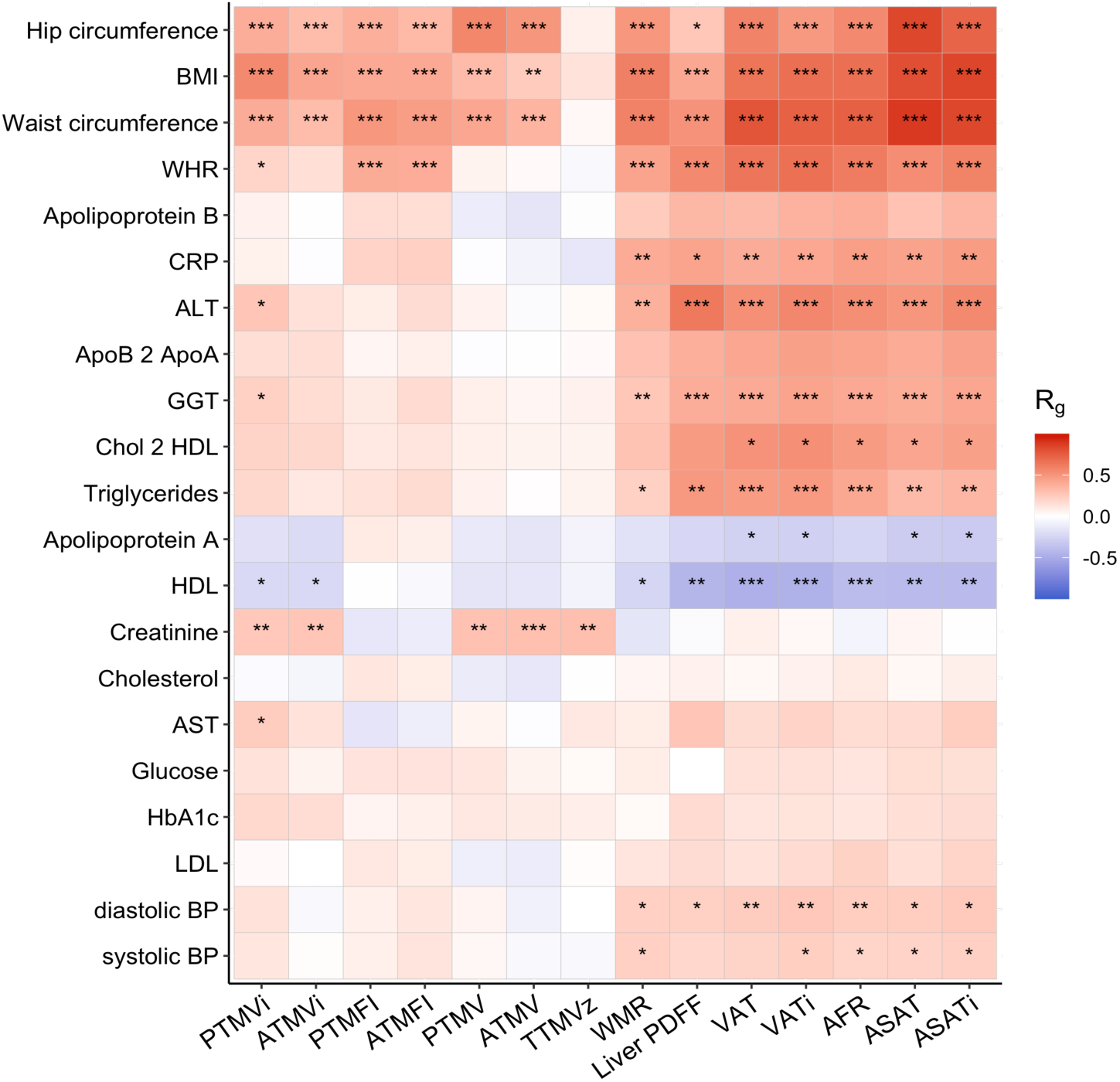
Genetic correlations of the MRI-derived body composition measures with standard anthropometrics and cardiometabolic measures. Abbreviations: BMI=body mass index, WHR=waist-hip ratio, CRP=C-reactive protein, ALT=alanine aminotransferase, GGT= gamma-glutamyl transferase, HDL=high-density lipoproteins, AST=aspartate aminotransferase, HbA1c=glycated hemoglobin, LDL=low-density lipoproteins, BP=blood pressure.

### Genetic correlation with cardiovascular, metabolic and mental disorders

Next, we analyzed the genetic overlap of the MRI-derived measures with medical conditions previously linked to cardiometabolic health, selecting recent GWAS with adequate power.^51–57^ As shown in Figure 5a, the strongest association across all measures was found for liver fat, with a genetic correlation of 0.48 (p=1.6×10^−22^) with type 2 diabetes. Coronary artery disease was found to have highly significant positive genetic correlations with visceral and subcutaneous adipose tissue. Overall, we found weak negative genetic correlations with muscle tissue measures and stronger positive genetic correlations with adipose tissue measures, with two exceptions; anorexia nervosa showed the opposite direction of correlation compared to the other conditions, and there was no discernible pattern for schizophrenia. Genetic correlations with the anthropometric and metabolic measures are provided in Figure 5b for comparison, indicating that the adipose tissue measures are as strong as or stronger correlated with these conditions than the conventional body anthropometrics.

**Figure 5.**
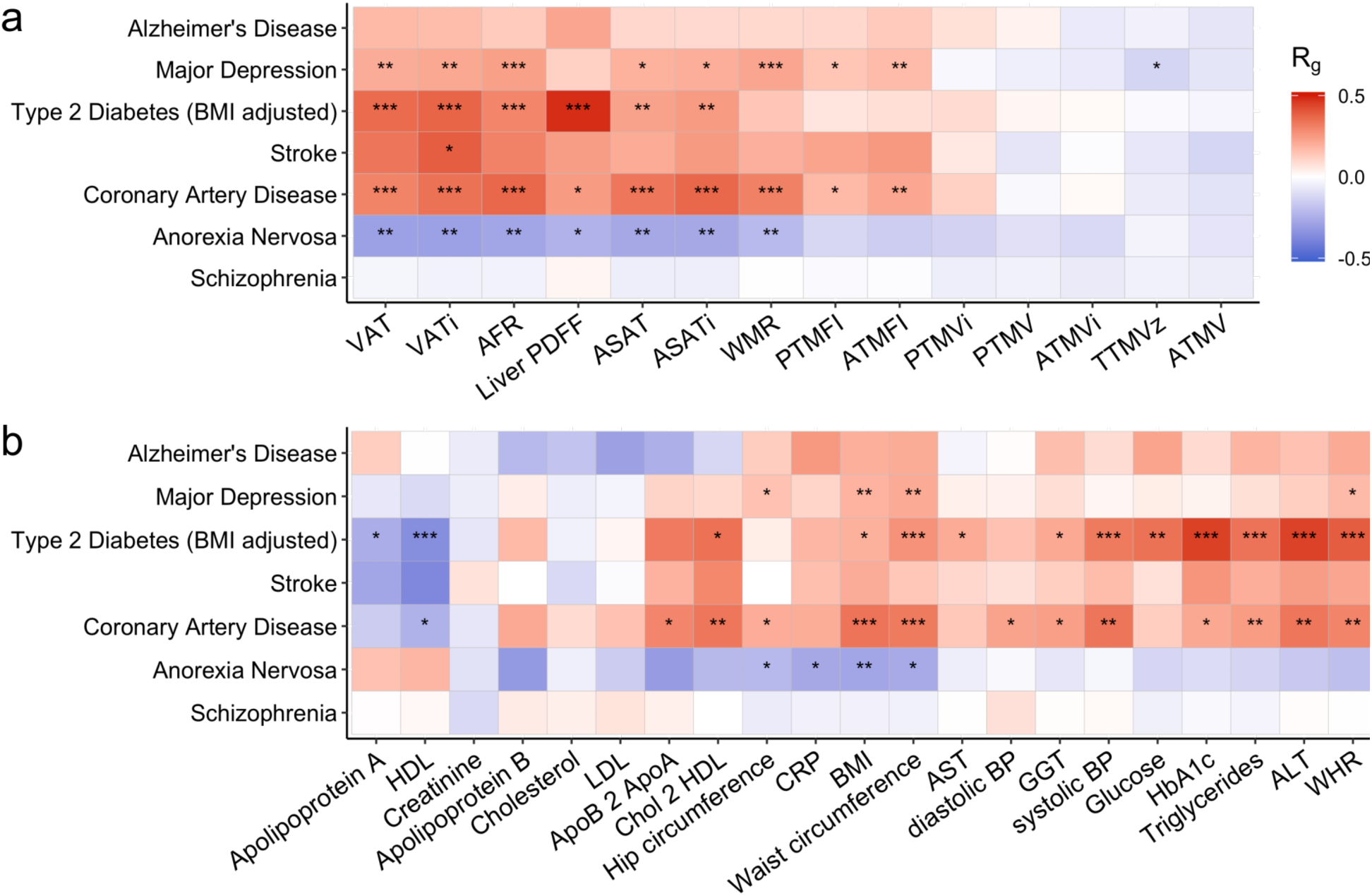
Genetic correlations of **a**) MRI-derived body composition measures, and **b**) anthropometric and metabolic measures (x-axis) with conditions linked to poor cardiometabolic health (y-axis). ***p=5×10^−9^, **p=5×10^−6^, *p=5×10^−4^.

## Discussion

Here, we reported results from a comprehensive, large-scale GWAS of MRI-derived measures of body composition. Joint analyses of measures of regional adipose and muscle tissue distributions revealed extensive genetic overlap and led to the identification of a large number of shared genetic risk loci across traits. We further showed genetic overlap with body anthropometrics and cardiometabolic measures as well as medical conditions linked to cardiometabolic health. Our findings illustrate how MRI-derived measures can be leveraged to improve our understanding of the biology underlying the metabolic system, identifying liver fat as a particularly promising measure, highlighting the integral role of steatosis and MAFLD in cardiometabolic health.

The genetic correlations of body composition measures with common medical conditions underlined that they may complement conventional measures to better understand cardiometabolic health. Liver fat showed a stronger genetic correlation with type 2 diabetes than conventional measures. The amount of liver fat and its genetic determinants may thus play a central role in type 2 diabetes development, and at a minimum robustly positions MAFLD onto the map of relevant comorbidities of type 2 diabetes alongside cardiovascular disease, kidney disease and diabetic retinopathy. Further, we found significant positive genetic correlations between coronary artery disease and visceral and subcutaneous adipose tissue, adding genetic evidence to the well-established relation between this disease, obesity, and body fat distribution.^58^

Liver fat also stood out from the other measures with regard to its genetic architecture. While all traits investigated were substantially heritable, the genetic discoverability of liver fat was much higher, with an oligogenic architecture as opposed to the polygenic architectures of the remaining traits and other complex biomedical measures.^45^ This was reflected in the GWAS yield, with a few highly significant loci coupled to lipid homeostasis explaining the majority of genetic variance for this measure. These loci should be scrutinized for the biological link between liver fat and cardiometabolic conditions,^59^ and may potentially point to fundamental processes that become dysregulated in these diseases. Indeed, all components of metabolic syndrome correlate with liver fat content.^60^ Evaluation of MAFLD risk has been recommended for any individual with metabolic syndrome and related morbidities (e.g. type 2 diabetes),^11,60^ and the large effects of these liver fat-associated loci even may suggest potential as features for individual risk stratification in MAFLD.^61,62^ These findings also attest to the accuracy and clinical relevance of MRI-derived measures of liver fat, and support the notion that MAFLD should be considered an integral component of obesity and metabolic syndrome and a key non-communicable disease.^11^

Another key finding was that the highest number of significant loci were found for muscle fat infiltration in the anterior and posterior thighs, two measures not previously genetically studied. Fatty infiltration of skeletal muscle reduces the muscle mass and strength,^63^ and has been implicated in aging and frailty.^64^ It has also been coupled to metabolic syndrome^65^ and cardiovascular mortality.^66^ Recent literature focused on liver disease and its progression have also highlighted the importance of muscle health.^67^ Muscle fat infiltration has been linked to higher comorbidity within MAFLD and decreased muscle fat infiltration has been correlated with improvement in steatohepatitis.^68,69^ Our findings suggest a strong genetic component to these associations, indicated by the large degree of shared genetic architecture with related diseases. Interestingly, fat accumulation in the muscle arises through specific pathways, including the intramyocellular accumulation of lipid,^63^ which is associated with insulin insensitivity and inflammation.^70^

The genetic correlations between the MRI-derived body composition measures indicate partly overlapping biological processes with some unique genetic determinants. The correlation structure further suggests that adipose tissue distribution is genetically largely independent from muscle tissue. However, it should be noted that global correlations underestimate overlap when a mixture of genetic effects in the same and opposing directions cancels each other out.^46^ Indeed, adipose and muscle tissue are known to have complex regulatory cross-talk, both releasing metabolism-regulating molecules to maintain a balanced weight-to-muscle ratio.^71^ The increased yield from the multivariate GWAS analysis, nearly doubling the number of unique loci discovered, is in line with the hypothesis of strong biological interplay and shared molecular mechanisms. The multivariate GWAS allowed for identifying loci that have distributed effects across the included body composition measures. These may help to explain the complexity of metabolic syndrome and the frequent comorbidity between diseases associated with body composition. Our findings that a substantial portion of the genetic determinants of these measures are related to the immune system fit with a large body of literature indicating that adipose tissue is an active metabolic and endocrine organ that secretes a host of pro- and anti-inflammatory factors, and with the characterization of obesity as a state of chronic low-grade inflammation.^72^ Thus, the current genetic findings can form the basis for functional follow-up studies to determine the molecular mechanisms involved in the complex relations between lipids and the immune system.

There was high genetic overlap between the sets of MRI-derived measures of body composition and the conventional measures of body anthropometrics and cardiometabolic health, indicating that they tag similar biological processes. The body anthropometrics were correlated with both muscle and adipose tissue, indicating little specificity, in line with the long-standing recognized limitations of these global measures that they fail to distinguish between specific body types that differ widely in risk for disease.^73^

Strengths of this study are the large number of whole-body MRI scans and the use of state-of-the-art, precise body composition measures, including multiple measures not previously investigated. With this, we were able to replicate loci reported earlier in smaller samples and with different measurement protocols.^35^ We further combined the study of individual measures with a multivariate approach to genetic discovery, allowing for greater GWAS hit yield and insight into the overall architecture of these complementary indicators of body composition and associated diseases. The findings allow for numerous follow-up investigations; further studies are needed to clarify the causal directions between the measures, and the role of putative moderators such as sex,^74^ age, and ethnicity.^75^

To conclude, the high prevalence of cardiometabolic diseases, combined with substantial morbidity and mortality, indicates a strong need for new therapeutic targets. While these diseases are often comorbid, they are treated separately, with this polypharmacy bringing along increased risk of adverse drug reactions.^4^ Genetic data is less subjected to reverse causation and confounding than environmental factors. Knowledge about shared and specific genetic determinants is therefore central to develop effective strategies that optimally treat the individual. We showed that accurate MRI-derived measures of liver and regional adipose and muscle tissue characteristics have strong genetic components, with shared influences that can be leveraged to boost discovery. As such, these findings have the potential to significantly enhance our understanding of body composition and related diseases, provide drug targets for MAFLD and related traits, and contribute to combatting a significant, increasing threat to public health.

## Acknowledgements

This work was partly performed on the TSD (Tjeneste for Sensitive Data) facilities, owned by the University of Oslo, operated and developed by the TSD service group at the University of Oslo, IT-Department (USIT). Computations were also performed on resources provided by UNINETT Sigma2 - the National Infrastructure for High-Performance Computing and Data Storage in Norway.

## Materials & Correspondence

The data incorporated in this work were gathered from public resources. The code is available via https://github.com/precimed/mostest (GPLv3 license), and GWAS summary statistics are uploaded to the GWAS catalog (https://www.ebi.ac.uk/gwas/). Correspondence and requests for materials should be addressed to

## Methods

### Participants

We made use of data from participants of the UKB population cohort, obtained from the data repository under accession number 27412. The composition, set-up, and data gathering protocols of the UKB have been extensively described elsewhere^76^. For the primary analyses, we selected White Europeans that had undergone the body MRI protocol, with available genetic and complete covariate data (N=34,036, mean age 64.5 years (SD=7.4), 51.5 % female). For the replication analyses, we made use of data from non-White Europeans (N=5081, mean age 63.0 years (SD=7.7), 52.9 % female).

### Data collection and pre-processing

Body and liver MRI scans were collected from three scanning sites throughout the United Kingdom, all with identical scanners and protocols. They were acquired on 1.5T Siemens MAGNETOM Aera scanners using a body dual-echo Dixon Vibe protocol and a single-slice multi-echo gradient Dixon acquisition, respectively. The UKB core neuroimaging team has published extensive information on the applied scanning protocols and procedures, which we refer to for more details.^77^ We acquired the data as processed by AMRA (Linköping, Sweden; https://www.amramedical.com), subsequently released by UKB. We bridged with UKB project accession #6569 to obtain early access to this data, which was then obtained from the UKB data repositories and stored locally at the secure computing cluster of the University of Oslo.

The AMRA body MRI processing included intensity inhomogeneity correction, non-rigid registration of atlases to acquired image volumes, quantification of fat and muscle composition using a voting scheme, and visual inspection for segmentation accuracy and manual adjustment.^78^ The liver MRI data were processed using a magnitude-based chemical shift technique with a 6-peak lipid model and then registered to the body MRI data and corrected for liver T2* to obtain a T1-weighted measure of liver proton density fat fraction (PDFF). AMRA implements manual quality control of the image/segmentation quality.

### Measurement protocols and definitions

We extracted a selection of body composition measures (Table 1; see also UKB online documentation (http://biobank.ctsu.ox.ac.uk/showcase/)). Specifically, we extracted the following measures of adipose tissue: visceral adipose tissue (VAT), defined as the adipose tissue within the abdominal cavity, and abdominal subcutaneous adipose tissue (ASAT), defined as the adipose tissue between the top of the femoral head and the top of T9. We also extracted measures of muscle fat infiltration (MFI) derived from the anterior and posterior thighs, averaged over both legs, and liver proton density fat fraction (PDFF). As measures of muscle tissue, we included anterior and posterior thigh fat-free muscle volume (ATMV and PTMV), and total thigh muscle volume, encompassing the gluteus, iliacus, adductors, hamstrings, quadriceps femoris, and sartorius, normalized to a z-score (TTMVz) that corrects for BMI, age, sex and height.^36^ We extracted two ratios from the UKB repository, namely weight-to-muscle ratio (WMR), defined as weight/TTMV, and abdominal fat ratio (AFR), which is (VAT+ASAT)/(VAT+ASAT+TTMV). Additionally, for VAT, ASAT, ATMV, and PTMV, we computed index measures by dividing these measures by the squared standing height in meters (e.g., ASATi is ASAT/height^2^). This is done since weight, adipose tissue, and lean tissue compartments scale to approximate height squared.

We subsequently regressed out age, sex, scanner site, and the first twenty genetic principal components from each measure. Following this, we applied rank-based inverse normal transformation^79^ to the residuals of each measure, leading to normally distributed measures as input for the GWAS.

For the secondary analyses, comparing the set of MRI-derived measures of body composition to measures of cardiometabolic health, we included 21 measures available in the UKB as listed in Table 2.

### GWAS procedure

We made use of the UKB v3 imputed data, which has undergone extensive quality control procedures as described by the UKB genetics team.^80^ After converting the BGEN format to PLINK binary format, we additionally carried out standard quality check procedures, including filtering out individuals with more than 10% missingness, SNPs with more than 5% missingness, and SNPs failing the Hardy-Weinberg equilibrium test at p=1*10^−9^. We further set a minor allele frequency threshold of 0.005, leaving 9,061,022 SNPs.

We carried out GWAS through the freely available MOSTest software (https://github.com/precimed/mostest). Details about the procedure and its extensive validation have been described previously.^47^ GWAS on each of the pre-residualized and normalized measures were carried out using the standard additive model of linear association between genotype vector, *g*_*j*_, and phenotype vector, *y*. Independent significant SNPs and genomic loci were identified in accordance with the PGC locus definition, as also used in FUMA SNP2GENE.^81^ First, we selected a subset of SNPs that passed genome-wide significance threshold 5×10^−8^, and used PLINK to perform a clumping procedure at LD r2=0.6 to identify the list of independent significant SNPs. Second, we clumped the list of independent significant SNPs at LD r2=0.1 threshold to identify lead SNPs. Third, we queried the reference panel for all candidate SNPs in LD r2 of 0.1 or higher with any lead SNPs. Further, for each lead SNP, its corresponding genomic loci were defined as a contiguous region of the lead SNPs’ chromosome, containing all candidate SNPs in r2=0.1 or higher LD with the lead SNP. Finally, adjacent genomic loci were merged if separated by less than 250 KB. Allele LD correlations were computed from EUR population of the 1000 genomes Phase 3 data. We made use of the Functional Mapping and Annotation of GWAS (FUMA) online platform (https://fuma.ctglab.nl/) to map significant SNPs from the MOSTest analyses to genes.

### MiXeR analysis

We applied a causal mixture model^43,44^ to estimate the percentage of variance explained by genome-wide significant SNPs as a function of sample size. For each SNP, *i*, MiXeR models its additive genetic effect of allele substitution, *β*_*i*_ as a point-normal mixture, 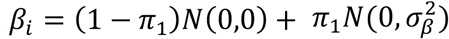, where *π*_1_ represents the proportion of non-null SNPs (‘polygenicity’) and 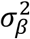 represents the variance of effect sizes of non-null SNPs (‘discoverability’). Then, for each SNP, *j*, MiXeR incorporates LD information and allele frequencies for 9,997,231 SNPs extracted from 1000 Genomes Phase3 data to estimate the expected probability distribution of the signed test statistic, 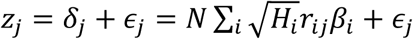, where *N* is the sample size, *H*_*i*_ indicates heterozygosity of i-th SNP, *r*_*ij*_ indicates an allelic correlation between i-th and j-th SNPs, and 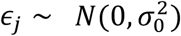 is the residual variance. Further, the three parameters, 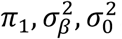, are fitted by direct maximization of the likelihood function. Fitting the univariate MiXeR model does not depend on the sign of *z*_*j*_, allowing us to calculate |*r*_*j*_| from MOSTest p-values. Finally, given the estimated parameters of the model, the power curve *S(N*) is then calculated from the posterior distribution p(*δ* | *z*_*j</sub>*,_*N*).

### Gene-set analyses

We carried out gene-based analyses using MAGMA v1.08 with default settings, which entails the application of a SNP-wide mean model and the use of the 1000 Genomes Phase 3 EUR reference panel. Gene-set analyses were done in a similar manner, restricting the sets under investigation to those that are part of the Gene Ontology biological processes subset (n=7522), as listed in the Molecular Signatures Database (MsigdB; c5.bp.v7.1).

## Notes

**Conflicts of interest** OAA has received speaker’s honorarium from Lundbeck and is a consultant to HealthLytix. JL, and ODL are employed by and stockholders in AMRA Medical, and RS was previously employed by AMRA medical. THK received consultancy fees from Intercept and Engitix and speaker fees from Novartis, Gilead and AlfaSigma. AMD is a Founder of and holds equity in CorTechs Labs, Inc, and serves on its Scientific Advisory Board. He is a member of the Scientific Advisory Board of Human Longevity, Inc. and receives funding through research agreements with General Electric Healthcare and Medtronic, Inc. The terms of these arrangements have been reviewed and approved by UCSD in accordance with its conflict of interest policies. All other authors report no potential conflicts of interest.

**Financial support** The authors were funded by the Research Council of Norway (276082, 213837, 223273, 204966/F20, 229129, 249795/F20, 225989, 248778, 249795, 298646, 300767), the South-Eastern Norway Regional Health Authority (2013-123, 2014-097, 2015-073, 2016-064, 2017-004, 2017- 112, 2019-101, 2020-060 (IES), 2022-080), Stiftelsen Kristian Gerhard Jebsen (SKGJ-MED-021), The European Research Council (ERC) under the European Union’s Horizon 2020 research and innovation programme (ERC Starting Grant, Grant agreement No. 802998), ERA-Net Cofund through the ERA PerMed project ‘IMPLEMENT’, and National Institutes of Health (R01MH100351, R01GM104400). This project has received funding from the European Union’s Horizon 2020 Research and Innovation Programme under Grant agreement No 847776847776 (CoMorMent).

### Competing Interest Statement

OAA has received speakers honorarium from Lundbeck and is a consultant to HealthLytix. JL, and ODL are employed by and stockholders in AMRA Medical, and RS was previously employed by AMRA medical. THK received consultancy fees from Intercept and Engitix and speaker fees from Novartis, Gilead and AlfaSigma. AMD is a Founder of and holds equity in CorTechs Labs, Inc, and serves on its Scientific Advisory Board. He is a member of the Scientific Advisory Board of Human Longevity, Inc. and receives funding through research agreements with General Electric Healthcare and Medtronic, Inc. The terms of these arrangements have been reviewed and approved by UCSD in accordance with its conflict of interest policies. All other authors report no potential conflicts of interest.

