## Supplements for "The role of liver fat in cardiometabolic diseases is highlighted by genome-wide association study of MRI-derived measures of body composition"

**Supplementary Information**

*List of discovered loci*

Supplementary Data 1 contains an overview of the individual loci discovered for each of the 14 primary univariate GWASs, including information on lead SNP, genomic location, significance, and mapped genes, as outputted by FUMA. Supplementary Figure 1 consists of the Manhattan plots, summarizing the significance of genetic variants across the genome for each of these univariate GWASs. Supplementary Data 2 contains the list of loci for the multivariate analysis of the primary set of MRI-derived measures. Last, Supplementary Data 3 lists the loci for the secondary multivariate analysis of the additional cardiometabolic measures.

*Significant genetic pathways and enrichment*

Please see Supplementary Data 4 for the top five most significant Gene Ontology biological processes pathways per primary measure (univariate and multivariate) as identified through MAGMA. Supplementary Data 5 lists significantly enriched pathways as determined through hypergeometric tests based on the set of mapped genes. Supplementary Figure 2 contains a visual representation of the Reactome analyses, coupling the set of genes identified through the primary MOSTest analysis to pathways, with Supplementary Data 6 containing the corresponding tabulated output, listing all significant pathways.

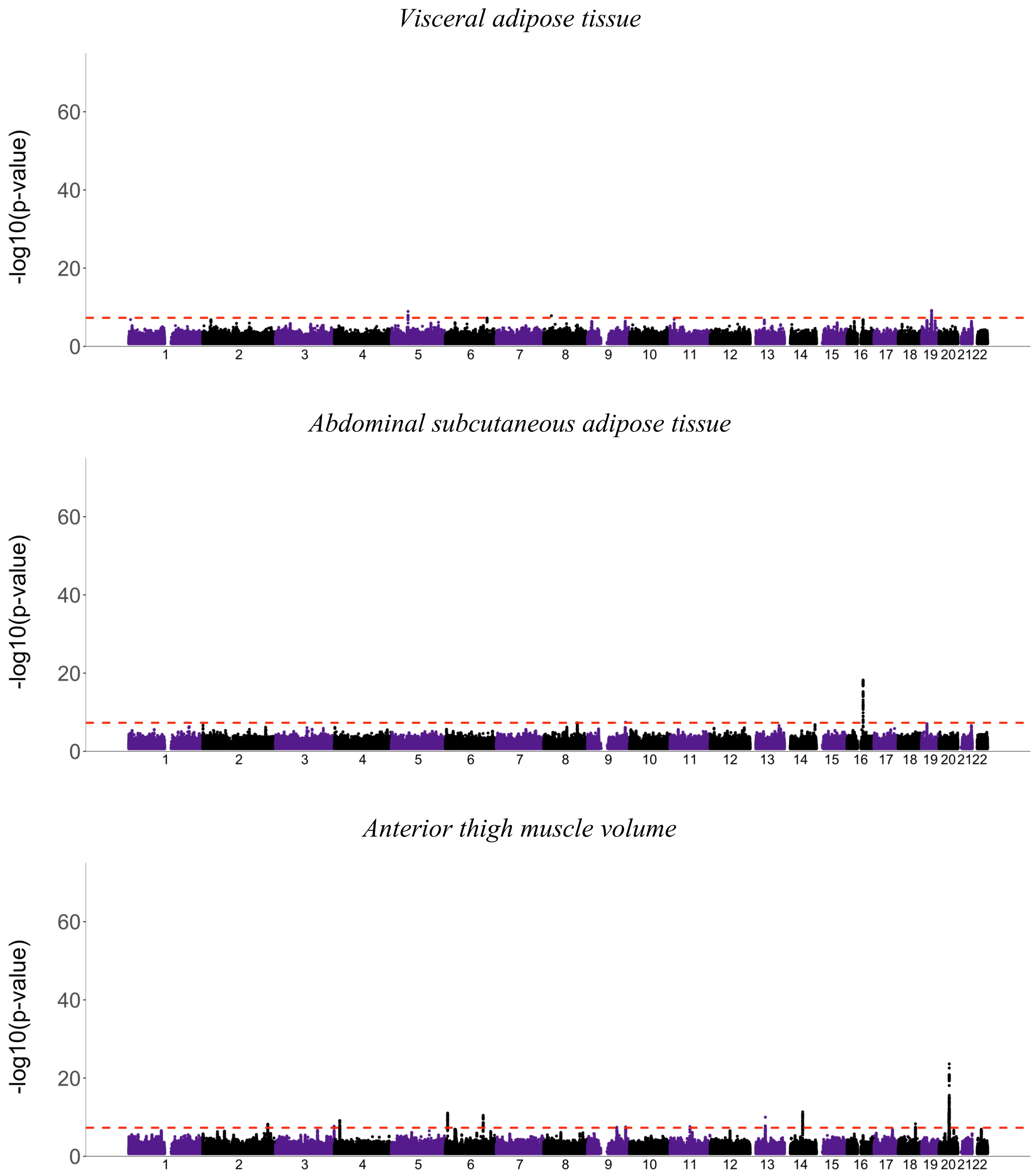
Visceral adipose tissue
-log10(p-value)
60
40
20
0
1 2 3 4 5 6 7 8 9 10 11 12 13 14 15 16 17 18 19 20 21 22
Abdominal subcutaneous adipose tissue
-log10(p-value)
60
40
20
0
1 2 3 4 5 6 7 8 9 10 11 12 13 14 15 16 17 18 19 20 21 22
Anterior thigh muscle volume
-log10(p-value)
60
40
20
0
1 2 3 4 5 6 7 8 9 10 11 12 13 14 15 16 17 18 19 20 21 22

*Posterior thigh muscle volume*

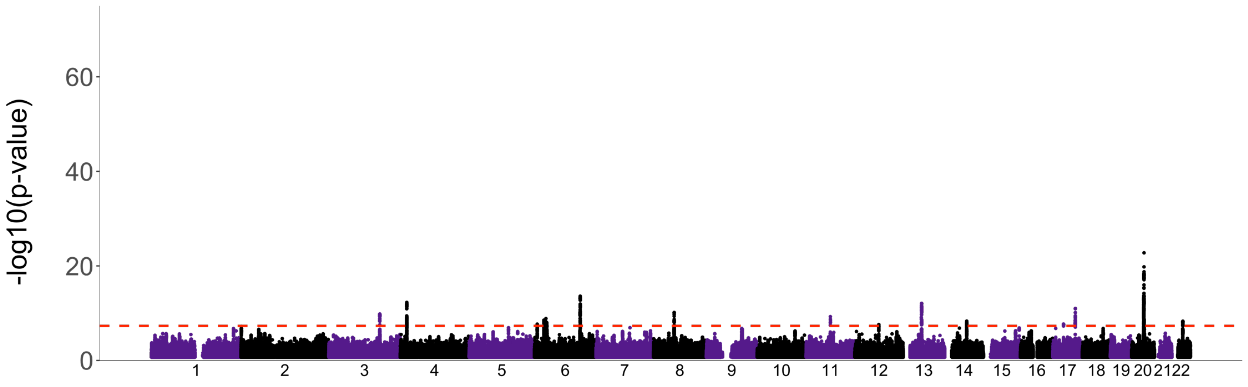


*Anterior thigh muscle fat infiltration (%)*

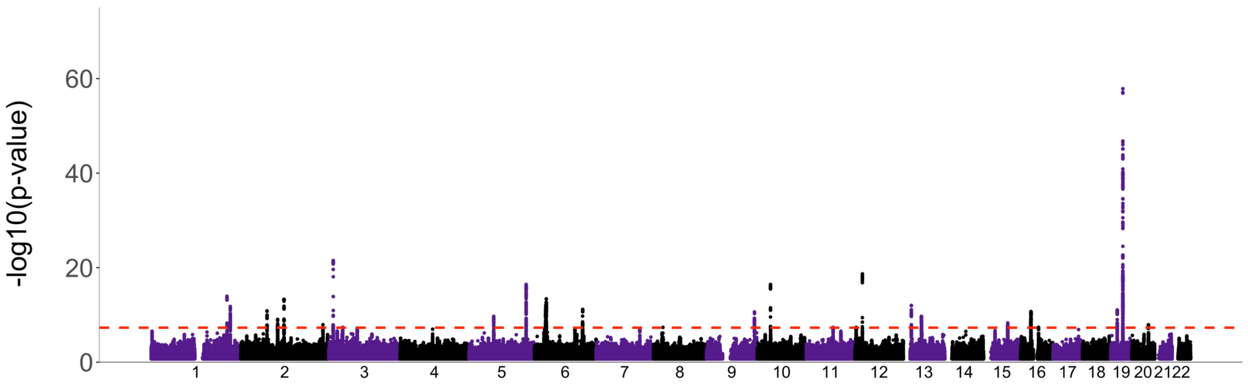


*Posterior thigh muscle fat infiltration (%)*

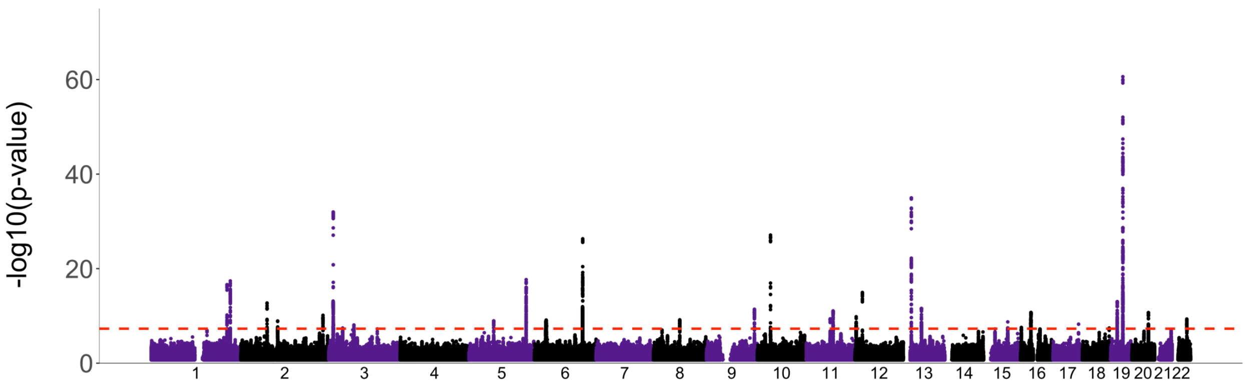

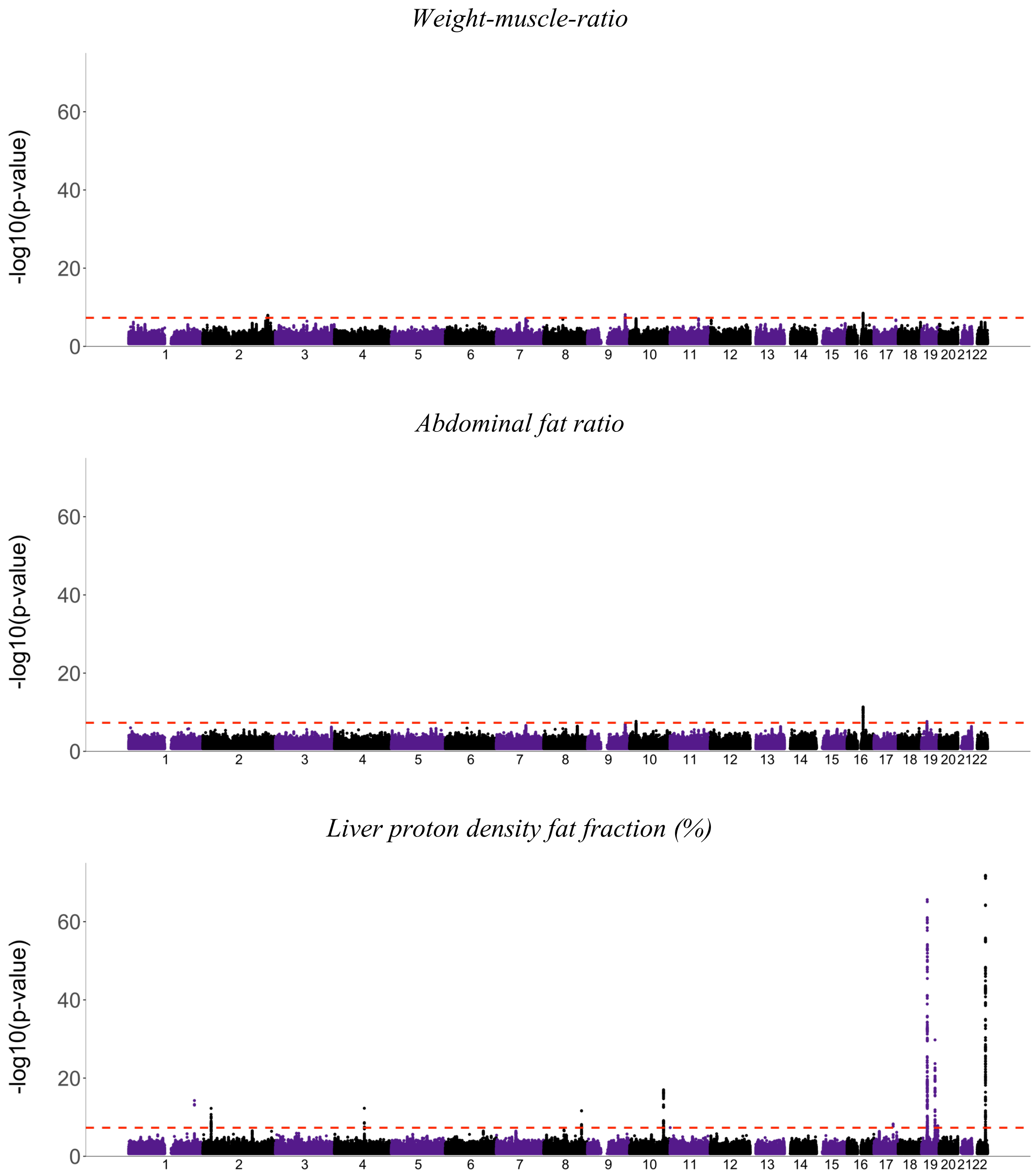
Weight-muscle-ratio
-log10(p-value)
Abdominal fat ratio
-log10(p-value)
Liver proton density fat fraction (%)
-log10(p-value)

*VAT/height*$^2$

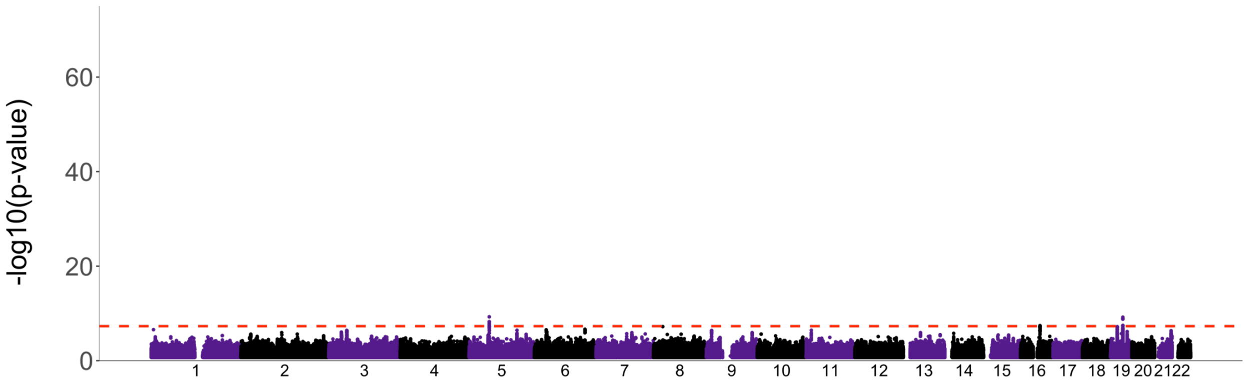


*ASAT/height*$^2$

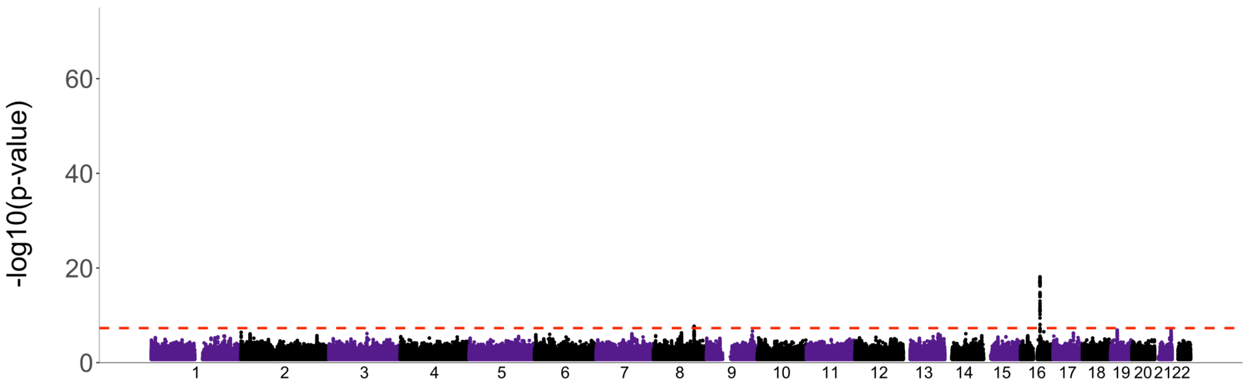


*ATMV/height*$^2$

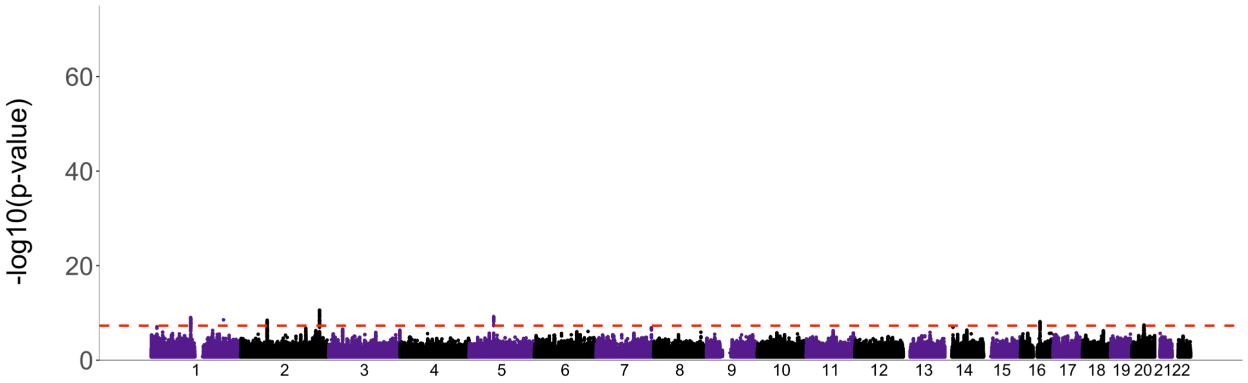

*PTMV/height$^2$*

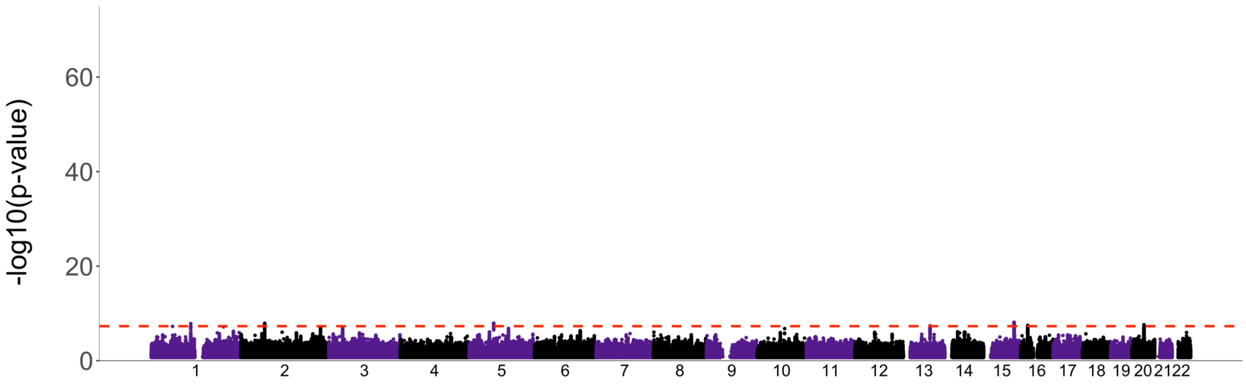


*Total thigh muscle volume (TTMVz; age, height, weight and sex corrected)*

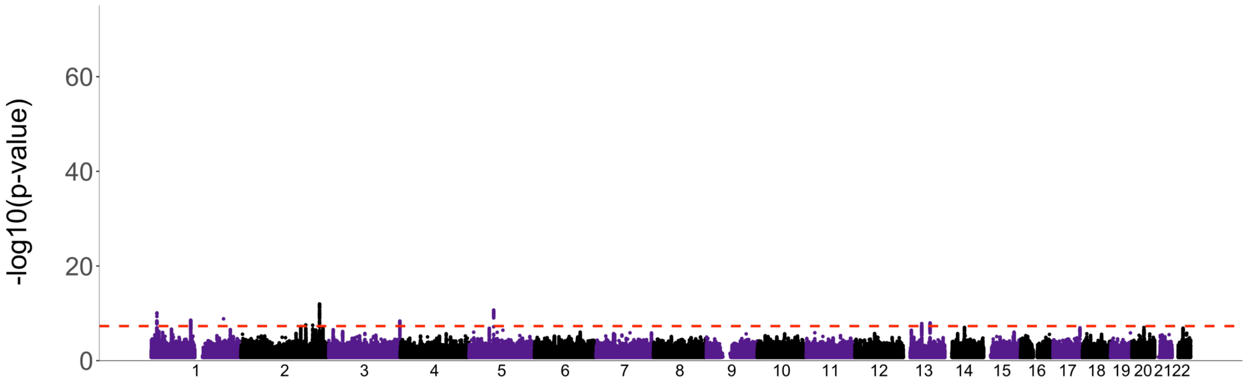


***Supplementary Figure 1. Manhattan plots for each of the MRI-derived measures of body composition, as indicated above each of the plots.*** *The x-axis indicates genomic position, ordered by chromosome, the y-axis indicates the -log10(p). The red horizontal dotted line indicates nominal GWAS significance threshold ($p=5*10^{-8}$).*

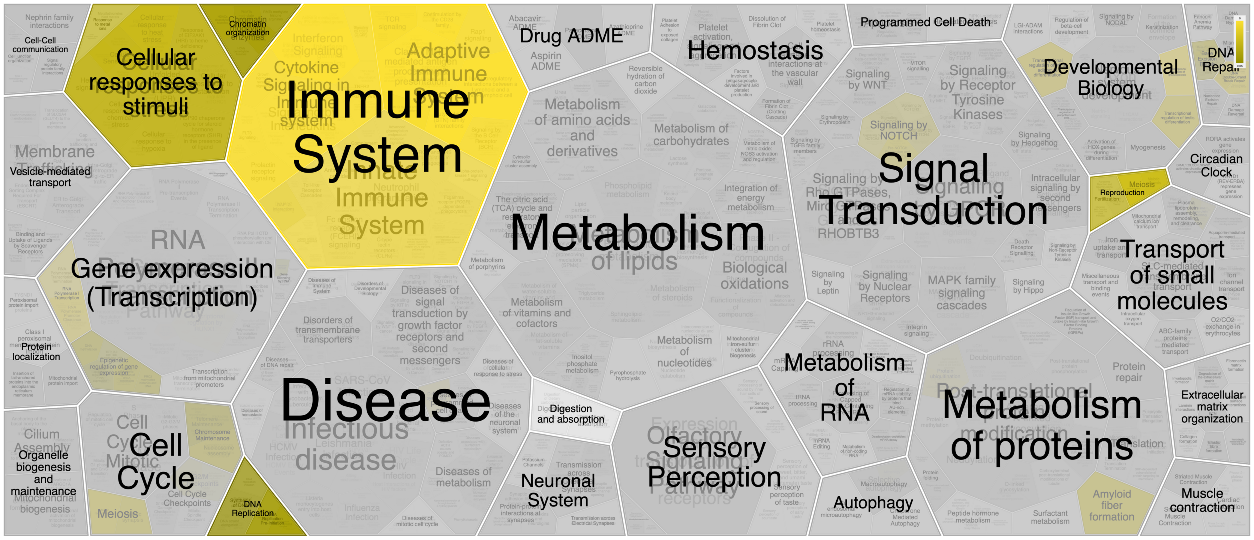


***Supplementary Figure 2. Visual representation of the output from coupling the MOSTest-identified set of genes to the Reactome database.*** *Brighter yellow indicates higher significance. Information on the specific pathways and their significance is provided in the Supplementary Data.*
